# Isolation and Genomic Analysis of Escherichia Phage AUBRB02: Implications for Phage Therapy in Lebanon

**DOI:** 10.1101/2024.06.17.599311

**Authors:** Tasnime Abdou Ahmad, Samar El Houjeiry, Antoine Abou Fayad, Souha Kanj, Ghassan Matar, Esber Saba

## Abstract

We obtained a new and unique Escherichia phage, AUBRB02, from sewage water in Beirut, Lebanon, as part of this research. AUBRB02 has an incubation period of around 45 minutes, a lysis period of about 10 minutes, and a burst size of around 30 plaque-forming units per cell. The phage exhibited strong biological stability over a pH range of 5.0–9.0 and temperatures ranging from 4°C to 60°C. AUBRB02 was found to have a genome size of 166,871 base pairs and a G+C content of 35.47% using whole-genome sequencing. A comparative analysis revealed that AUBRB02, a newly found phage, shares 93% intergenomic similarity to closest relative in refseq. Functional annotation revealed the presence of 10 tRNA and 262 coding sequences, out of which 123 are categorized as putative proteins. These results indicate that AUBRB02 is a highly infectious virus that belongs to the *Tequatrovirus* genus. This study is significant reference information that can be used in the development of phage therapy.

**IMPORTANCE:** *Escherichia coli*, a gram-negative bacterium, is a widely distributed pathogen in the natural environment and a frequent cause of illnesses. The extensive utilization of antibiotics has resulted in a rise of clinically resistant strains, posing a substantial obstacle to antimicrobial therapy. This urgent circumstance highlights the necessity for antibiotic substitutes to combat *E. coli* infections. In this context, we introduce AUBRB02, a novel Escherichia phage isolated from an untreated sewage source in Beirut. Our findings indicate that AUBRB02 is highly lytic, stable against extreme culturing conditions, and has a biofilm elimination capability.

## INTRODUCTION

*Escherichia coli* is a gram-negative rod-shaped bacterium that belongs to the Enterobacteriaceae family and is the most prevalent commensal inhabitant of the gastrointestinal tracts of humans and warm-blooded animals, while rarely causing disease to their host. It is however considered one of the most common human and animal opportunistic pathogens, owing to its ability to survive both aerobic and anaerobic conditions along with its wide availability [1].

Shortly after the discovery and release of the first antibiotic - penicillin, [2], penicillin-resistant Staphylococcus aureus strains emerged, paving the way to spread of antibacterial resistance, a current worldwide public health problem [3]. Several factors contribute to the origin of antibacterial resistance, such as conjugation, transduction, transformation [4–6], acquired genetic mutations [7], intrinsic natural resistance [8], etc. Lebanon shares along with other Eastern Mediterranean countries (low-income countries) a prevalence of resistance higher than several European countries, due to misuse of antibiotic in animal agriculture [9] that spread antibiotic resistant to humans [10], and inadequate wastewater treatment and disposal [11]. The inadequate wastewater treatment of hospitals, industrial and farm sewage lead to an increase in the rates of antibiotic resistance in *E. coli*. According to a study conducted at the American University of Beirut, 5 out of 8 *E. coli* isolated from patients prostate glands produced extended spectrum beta-lactamases (ESBL) [12]. Another study conducted at the same hospital, showed that 40 % of the *E. coli* isolated from ventilator-associated pneumonia patients produced ESBL [13]. A study conducted by *Dagher LA et al* in 2021 to assess the water quality in 14 different rivers water, detected the presence *E. coli* in 95.5% of the samples, respectively. Out of the isolated samples, 45.8% were classified as multi-drug resistant (MDR) [14]. In addition, a study conducted on 5 major rivers in Lebanon, showed major dissemination of MDR *E. coli* and *Klebsiella pneumoniae* isolates, among the collected samples [15]. This necessitated the production of new antibiotics, to target AMR pathogens particularly *E. coli* (categorized as critical pathogen by the World Health Organization in 2017) [16]. However, the discovery of new drugs is laborious and may take up to years, which made it difficult to keep up with the emergence and spread of AMR [17], which highlights the need for alternatives to antibiotics, particularly natural alternative, such as “bacteriophages”.

Bacteriophages are filterable viruses that infect or kill bacterial cells without causing any harm to host cells [18]. They are the most abundant and diverse biological entities on the planet, with each differing in its morphology, shape, and size [19]. They consist of an icosahedral protein capsid surrounding their genomic material, whether DNA or RNA single stranded or double stranded [20]. Bacteriophages can undergo five different life cycles, lytic, lysogenic, pseudo-lysogenic, chronic, and cryptic life cycle [21, 22]. However, and for the sake of this study we will focus only on the lytic phages that multiply using the host’s machinery (attachment, penetration, biosynthesis, maturation, assembly, and release), yielding bacterial cell death [23]. Unlike antibiotics, bacteriophage during bacterial-killing process is capable of increasing their number, through auto-dosing upon host identification and infection [24, 25]. To date several bacteriophages have been isolated and characterized against *E. coli*, and were even subjected to animal trails, including Ib_pec2, CS EPEC, BL EPEC, BI EPEC, CI EPEC, BL EHEC, BI EHEC, vB_EcoM_ESCO8, vB_EcoM_ESCO32, vB_EcoM_ESCO56, vB_EcoM_ESCO9, MLP 1, MLP 2, MLP 3, BC-S1, ETEC-S3, EHEC-S4, BC-S2, myPSH1131 [26–31]. Given the above, this study aims to isolate and characterize bacteriophages with lytic activity against multi-drug resistant *Escherichia coli* from Ramlet El-Bayda sewage water.

## RESULTS

### Isolation of Escherichia coli Phage

AUBRB02, was isolated using *E. coli* ATCC 25922 from a sewage sample collected at Ramlet El-Bayda. A clear, circular-shaped plaque was observed on the DLA plate and enriched on the same host at least 3 times. Phage AUBRB02 formed a clear plaque. This small circular zone, where the phage lysed the surrounding bacterial cells, highlights potential lytic activity of AUBRB02 (Figure 1).

**Figure 1:**
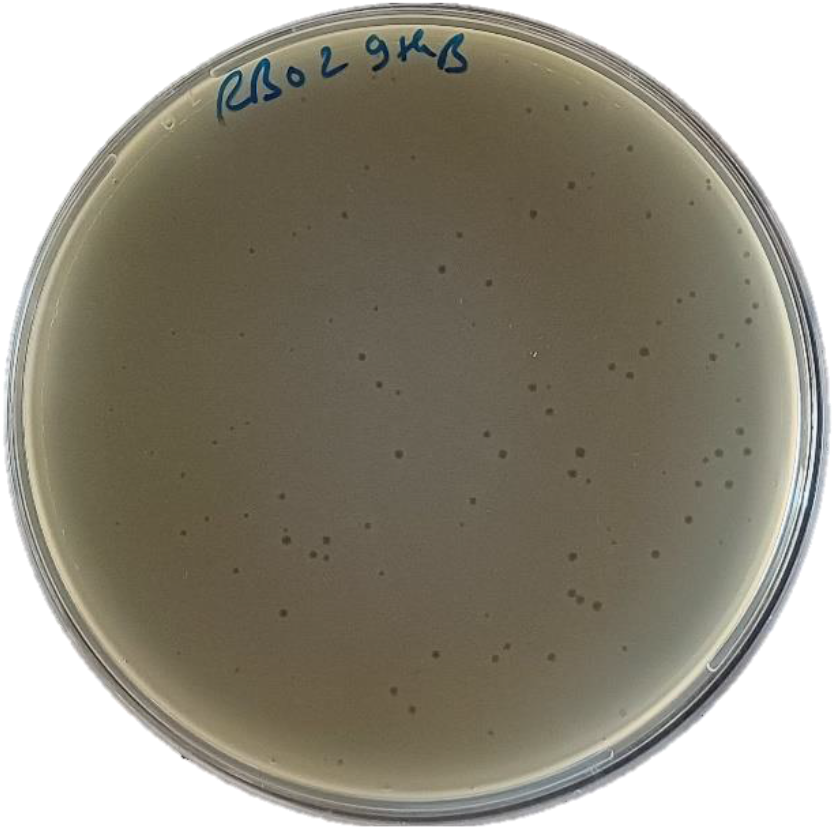
Plaque Formation of *Escherichia* Phage AUBRB02 on *E. coli* ATCC25922 Lawn.

### Bacteriophage host range determination

We tested the host range of *Escherichia* phage AUBRB02 against 18 *E. coli* strains, with susceptible strains highlighted in red in Figure 2. These strains, primarily isolated from urine, sputum, DTA and perianal samples. The phylogenetic tree indicates genetic diversity among the tested strains. Susceptible strains are dispersed across different clusters in the phylogenetic tree, indicating that phage AUBRB02 can infect genetically diverse *E. coli* strains regardless of their phylogenetic relatedness, suggesting a broad host range. The antibiotic resistance profiles are displayed in a heatmap, showing resistance (R), susceptibility (S), and intermediate (I) responses to 18 antibiotics across various classes, including β-lactams, aminoglycosides, quinolones, polymyxins, tetracyclines, glycylcyclines, and others. The data reveals high resistance levels among many strains, particularly to CAZ, CIP, TE, and SXT. Notably, phage-susceptible strains (marked in red) exhibit varying levels of resistance but tend to show less resistance compared to non-susceptible strains. For instance, *E. coli* 174, 179, 178, 180, 177, and 182, despite their susceptibility to phage AUBRB02, maintain susceptibility to a broader range of antibiotics compared to more resistant strains.

**Figure 2:**
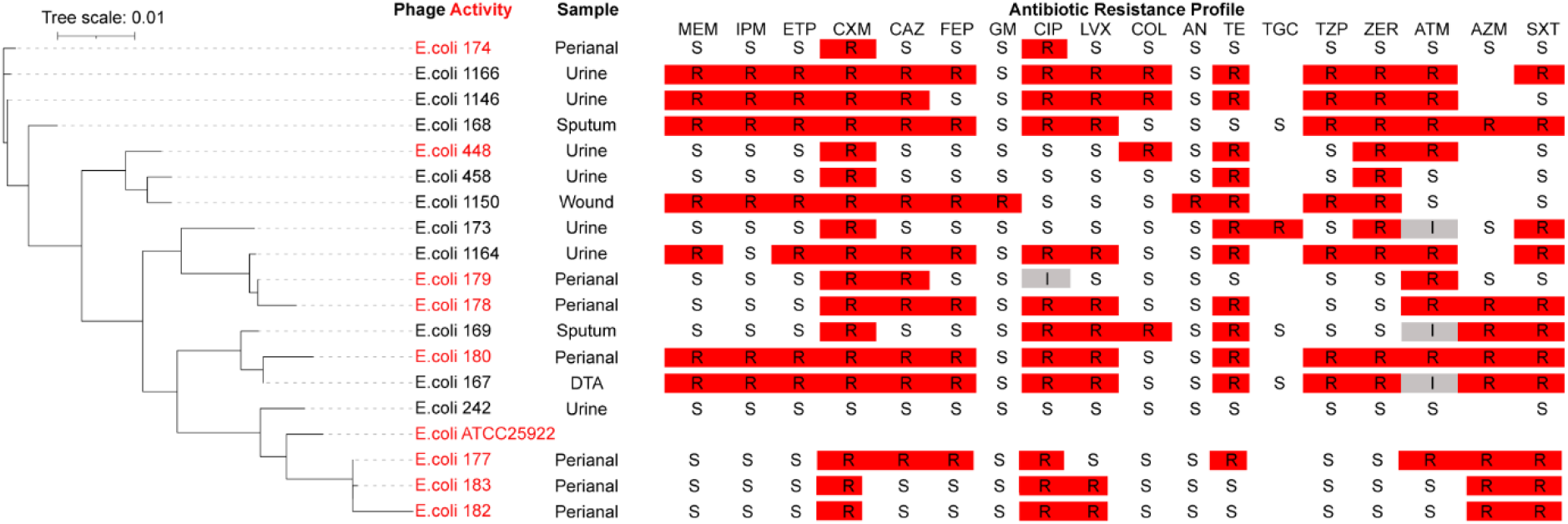
Host Range of *Escherichia* Phage AUBRB02 and Antibiotic Resistance Profiles of ***E. coli* Strains**. The phylogenetic tree on the left with a tree scale of 0.01 indicating genetic distance. Strains susceptible to *Escherichia* phage AUBRB02 are highlighted in red. The right section presents the antibiotic resistance profiles of the *E. coli* strains against 18 antibiotics. Red squares denote resistance (R), white squares denote susceptibility (S), and grey squares denote intermediate resistance (I).

### Bacteriophages one-step growth and adsorption rate

The adsorption rate and one-step growth curve of *Escherichia* phage AUBRB02 on the host strain *E. coli* ATCC 25922 were examined to understand the phage’s interaction with its host. The adsorption rate data revealed that within the first 5 minutes, approximately 80% of the phages adsorbed to the host cells, followed by a plateau phase from 10 to 25 minutes (Figure 3, A). This rapid initial adsorption phase suggests a high affinity of the phage for the bacterial cells. The one-step growth curve analysis showed an initial latent phase of about 45 minutes, during which the phage particles are not yet released. This was followed by a burst phase, with the phage titer peaking at around 400 plaque-forming units per milliliter (PFU/ml) after 60 minutes. This peak indicates the release of new phage particles from lysed host cells, with a calculated burst size of approximately 250 virions per bacterial cell. These findings highlight the efficiency of phage AUBRB02 in infecting and lysing *E. coli* ATCC 25922 cells (Figure 3, B).

**Figure 3:**
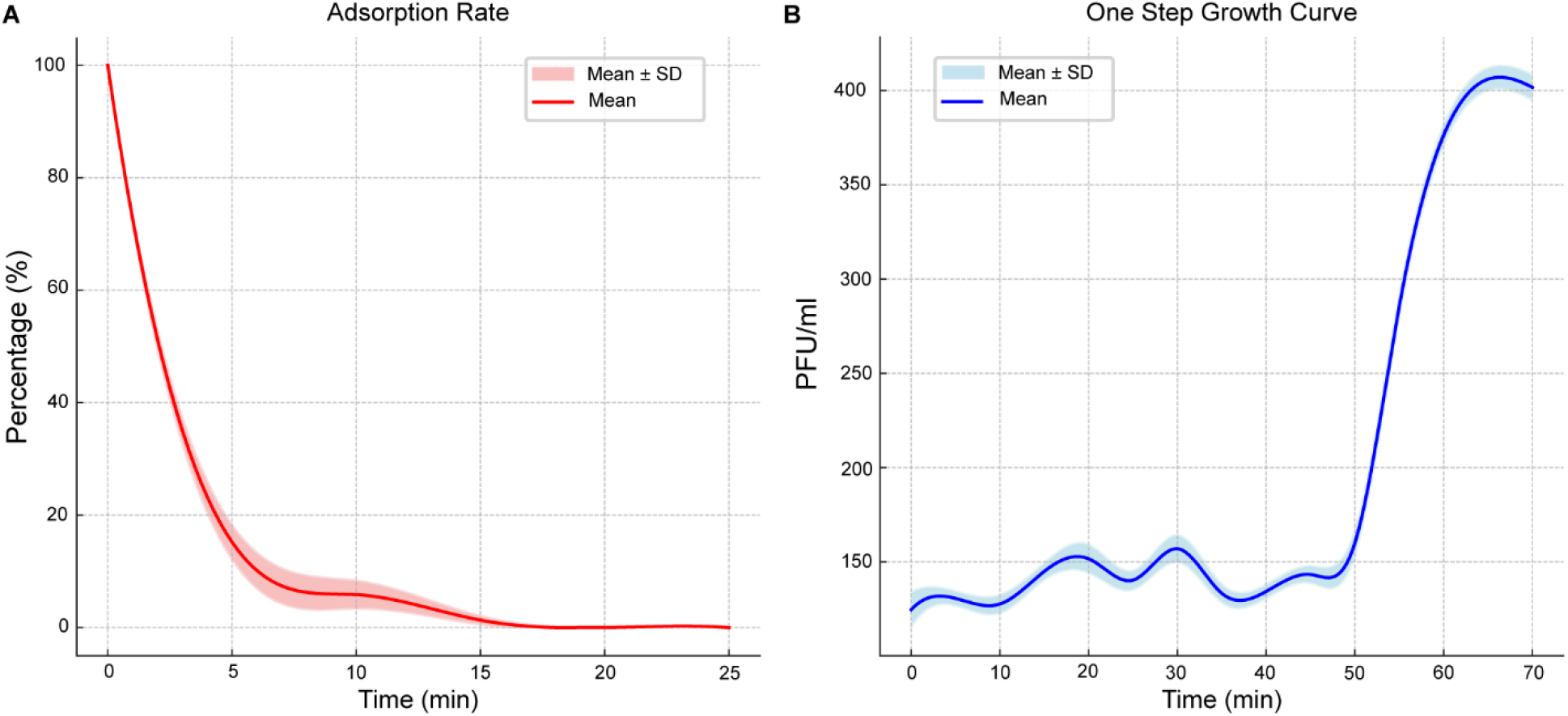
Adsorption Rate and One-Step Growth Curve of *Escherichia* Phage AUBRB02. (A) Adsorption Rate: The percentage of free phage particles not adsorbed to *E. col*i cells over 25 minutes is shown. (B) One-Step Growth Curve: The phage titer in PFU/ml over time. Shaded area represents Mean ± SD.

### Bacteriolytic activity

The bacteriolytic activity of *Escherichia* phage AUBRB02 was determined at different MOIs (10, 1, 0.1, and 0.01) against *E. coli* ATCC 25922. As shown in Figure 4, the optical density at 600 nm (OD600) was measured over time. The positive control (phage-free bacterial culture) showed a continuous increase in absorbance over 13 hours, while the negative control (media without bacteria) remained constant. For the treated wells, the absorbance initially increased to a maximum in an MOI-dependent manner within 2-3 hours, followed by a rapid decline to levels resembling the negative control within 4 hours. At higher MOIs (10 and 1), a rapid decline in OD600 was observed within the first 3 hours, indicating efficient bacterial lysis. Lower MOIs (0.1 and 0.01) showed a slower decline and gradual reduction in bacterial density, demonstrating less aggressive but still effective lytic activity. This data confirms the effectiveness of phage AUBRB02 in lysing *E. coli* ATCC 25922 across a range of MOIs, with higher MOIs resulting in more rapid bacterial lysis.

**Figure 4:**
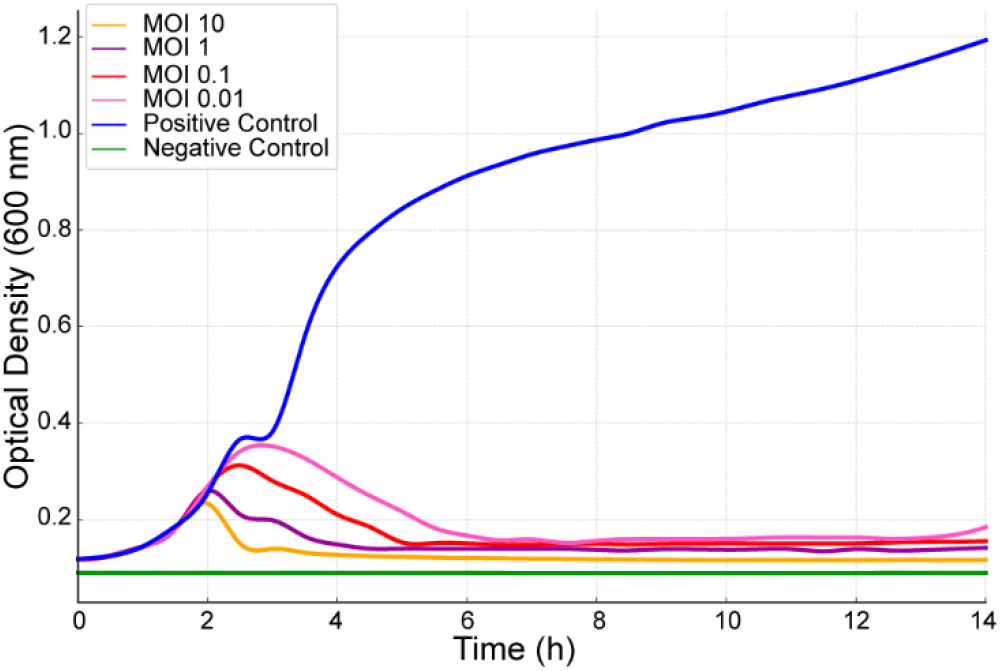
Impact of Different MOIs on the Growth of *E. coli* ATCC 25922 When Infected with *Escherichia* Phage AUBRB02. The graph shows the optical density at 600 nm (OD600) over time for *E. coli* ATCC 25922 treated with *Escherichia* phage AUBRB02 at MOIs of 10, 1, 0.1, and 0.01 (orange, pink, red, and yellow lines). The blue line represents the positive control (phage-free bacteria), and the green line is the negative control (media only).

### pH and thermal stability

The stability of *Escherichia* phage AUBRB02 was assessed under varying temperature and pH conditions, revealing important insights into its robustness (Figure 5). The phage titer, represented in 10^10^ PFU/ml, remained high (approximately 20×10^10^ PFU/ml) across temperatures ranging from 10°C to 60°C, indicating strong stability within this range. However, a sharp decline in titer was observed above 60°C, with the phage becoming completely inactivated at 80°C. Similarly, pH stability tests showed that phage AUBRB02 maintained a high titer across a pH range of 4 to 10, with peak stability occurring around neutral pH (7 to 8). Outside this range, particularly below pH 4 and above pH 10, the phage titer dropped significantly, indicating reduced stability in highly acidic or basic conditions. These findings highlight that while *Escherichia* phage AUBRB02 is robust across a range of moderate environmental conditions, it is sensitive to extreme temperatures and pH levels, which is crucial for its storage and application in various settings.

**Figure 5:**
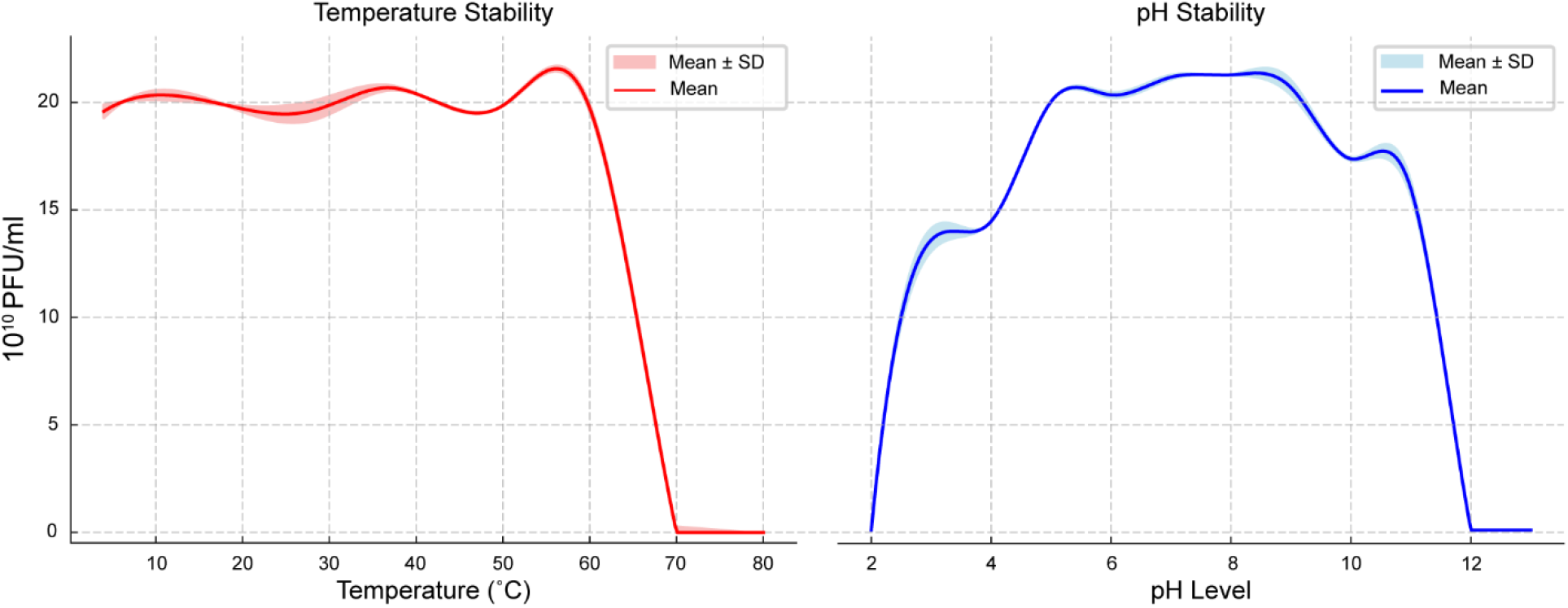
Temperature and pH Stability of *Escherichia* Phage AUBRB02. The phage titer (10^10^ PFU/ml) remains stable between 10°C and 60°C, but drops sharply above 60°C, with complete inactivation at 80°C. The phage shows stability across a pH range of 4 to 10, with peak stability around pH 7 to 8. Titer decreases significantly below pH 4 and above pH 10.

### Inhibition of biofilm formation

The efficacy of *Escherichia* phage AUBRB02 in reducing biofilm post-formation was assessed. Upon treatment with *Escherichia* phage AUBRB02, there was a substantial reduction in biofilm, as evidenced by significantly lower OD values compared to the positive biofilm control, demonstrating the phage’s effectiveness in biofilm disruption. These results underscore the specificity and potency of *Escherichia* phage AUBRB02 in targeting and reducing *E. coli* biofilm post-formation, highlighting its potential application in combating biofilm-associated infections (Figure 6).

**Figure 6:**
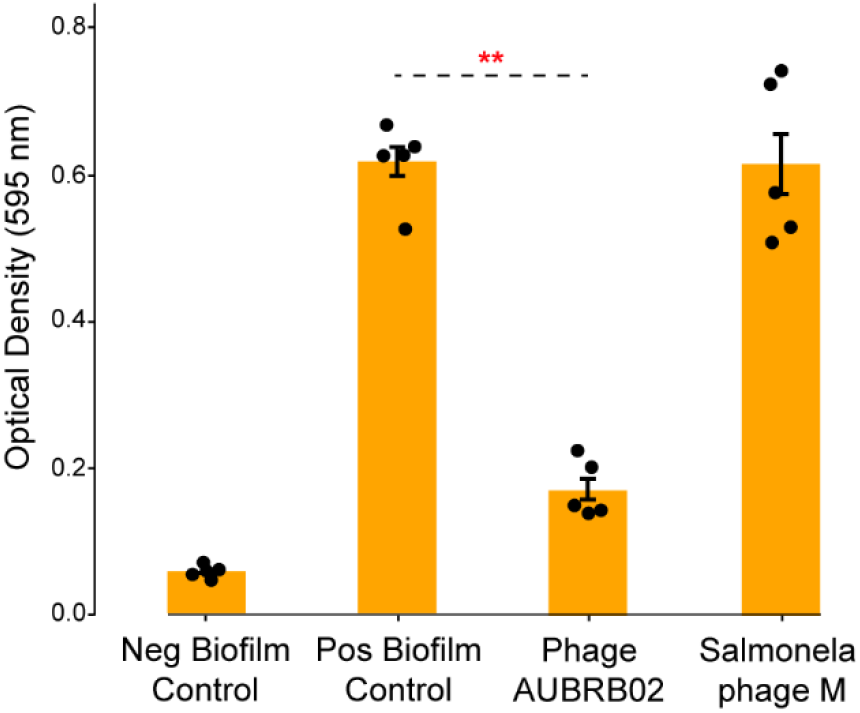
Inhibition of Biofilm Post-Formation by Phage Treatment. *E. coli* ATCC 25922 biofilms were treated with AUBRB 02 phage (10^9^ PFU/ml) in a 96-well plate. After 24 hours, biofilms were stained with crystal violet, and optical density at 595 nm (OD595) was measured. The negative biofilm control, positive biofilm control, and Salmonella phage M control are shown for comparison. Data are mean ± SD. Statistical significance was determined by one-way ANOVA, with **P < 0.01 between the positive control and phage AUBRB02.

### Whole genome sequencing and bioinformatic analysis

Phage AUBRB02 has a circular double-stranded DNA with a 166871-bp-long genome and a GC content of 35.47%. The whole genome sequence of AUBRB02 was analyzed using BLASTn from the NCBI non-redundant DNA database. Comparative analyses conducted demonstrated that AUBRB02 clustered with *Straboviridae* phages. These phages alignment demonstrated that they had the same collinear genome arrangement (Figure 7, C). The VIRIDIC (Virus Intergenomic Distance Calculator) analysis revealed a 93% intergenomic similarity between the compared phage genomes. Thus, indicating that the genome sequence of AUBRB02 was relatively new. The detailed intergenomic similarity matrix is presented in supplementary data (Figure S1). Functional annotation using PROKKA revealed that AUBRB02 contained 262 CDSs and 10 tRNAs (Figure 7, A). The annotated proteins from the phage include a variety of functional families. Among the enzymes identified are DNA topoisomerase II, DNA primase, RNA ligase, and several helicases and polymerases. Regulatory proteins are well-represented, including transcriptional regulators such as FmdB-like and MotB-like regulators, sigma factors, and anti-sigma factors. Structural proteins encompass virion structural proteins, baseplate wedge subunits, tail fibers, and head scaffolding proteins. Numerous hypothetical proteins are present, reflecting uncharacterized or putative functions. Transport and membrane-associated proteins include RIIA and RIIB lysis inhibitors and holins. Additionally, the annotation reveals several tRNAs and other binding proteins, such as RNA polymerase binding proteins and internal virion proteins. Miscellaneous proteins include antitoxins, thioredoxins, ribonucleotide reductases, and several enzymes involved in nucleotide metabolism. Further analysis of the complete genome using phage scope and manual inspection revealed the absence of genes associated with temperate or lysogenic life cycles, and virulent- and antibiotic-related genes. This indicates that AUBRB02 has a strictly lytic life cycle and does not carry genes conferring lysogeny or antibiotic resistance.

**Figure 7:**
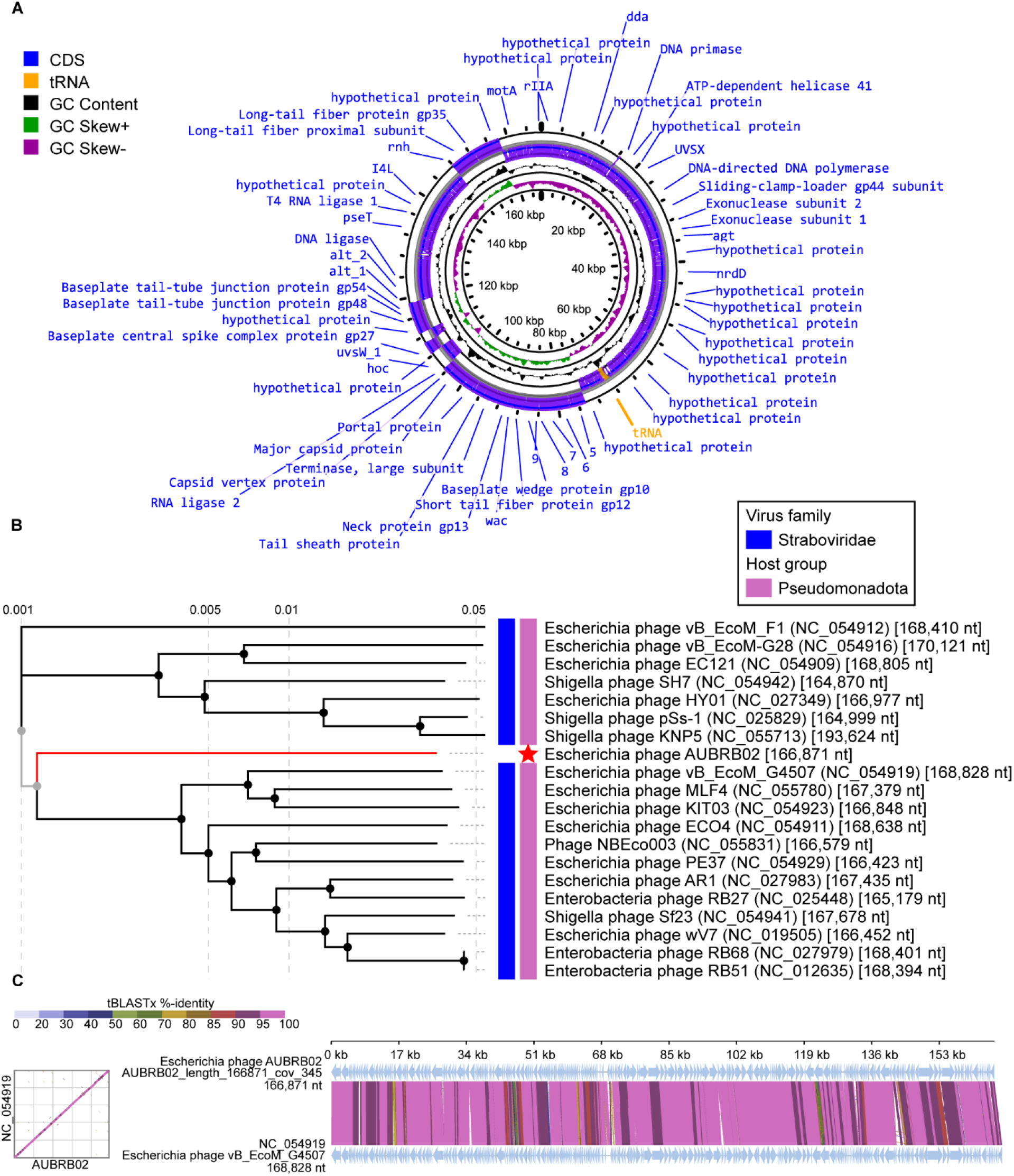
Genomic Analysis of *Escherichia* Phage AUBRB02. (A) Circular genome map showing coding sequences (CDS in blue), tRNAs (orange), GC content (black circle), and GC skew (green for positive, purple for negative). Key structural and functional genes are annotated. (B) Phylogenetic tree based on tBLASTx, with AUBRB02 (red star) clustered within the *Straboviridae* family, showing close relationships with other *Pseudomonadota* phages. (C) Dot esplot and synteny analysis demonstrating high sequence similarity between AUBRB02 and *Escherichia* phage vB_EcoM_G4507, indicating conserved genomic regions.

## DISCUSSION

In the current study, the phage isolated against *E. coli* ATCC 25922 was isolated from Ramlet El-Bayda untreated sewage source.

The ideal phage for phage therapy is a phage that targets a single bacterial species while targeting many if not all strains belonging to the targeted species [46]. The lysis effect (34 %) revealed that phage AUBRB 02 has the potential to be a candidate for phage therapy. Generally, phage receptor binding protein can interact with bacterial strains harboring the same bacterial cell receptor as that of the host strain [47]. Broad spectrum host range phage is essential for phage therapy as it can target several bacterial strains, resembling broad spectrum antibiotic in effect [48]. Nevertheless, bacteria have developed certain resistance strategies to interfere with phage-bacterial infection process. One of the most common limitation of phage therapy is adsorption resistance, where bacterium cell losses or hide bacterial binding receptors, thus restraining the phage and bacterial interaction [49].

As shown in Figure 3, AUBRB 02 recorded a latent period of 45 min accompanied by a 279 PFU/ml burst size. Short latent period and large burst size are essential features for highly effective phages serving as therapeutic agents [50, 51]. Previously, the burst size of the phages ϕAPCEc01, ϕAPCEc02 and ϕAPCEc03 belonging to *Myoviridae* and *Siphoviridae* was 90, 30 and 47 PFU/ml with a latent period of 60 min, 40 min and 40 min [52]. Another phage, *Escherichia coli* bacteriophage Ω8, recorded a latent period of 19 min and burst size of 130 PFU/ml [53]. In addition, more than 99% of the phages adsorbed and infected the bacterial cells after 15 min of incubation. Therefore, the observed results indicate that AUBRB 02 could be rapidly absorbed into the host cells, and latent period and burst size reflects the lytic effectiveness of AUBRB 02 towards *E. coli* ATCC 25922.

Stability of phages against a wide range of pH’s and temperatures is a major advantage for phage therapy. As shown in Figure 5, incubation at 70°C yielded inactive phages which aligns with the results reported in a previous study by Yamaki et al; where the phage’s nucleic acid and proteins denatured upon exposure to high temperatures [54]. Furthermore, AUBRB 02 incubation at pH 11 yielded an inactive phage, which might be due to phage capsid protein dissociation owing to high concentrations of hydrogen and hydroxyl ion in the solution [55].

One of the most common broad bacterial defense mechanisms is the formation of a biofilm matrix, which can trap the phage upon entry, therefore reducing the phage RBP interaction with their corresponding bacterial receptor [56]. Nonetheless, various studies have been reported targeting phages with parasitic activity against *E. coli* species, with these phages displaying biofilm elimination capacities [57–59]. In this study, we examined the ability of AUBRB 02 phage in eliminating mature biofilm using *E. coli* ATCC 25922, which has been used as a positive control when testing *E. coli* strains biofilm formation capabilities [60]. AUBRB 02 showed an excellent effect in reducing mature biofilm (biofilm post-formation). The annotated phage genome contains proteins that may play critical roles in biofilm degradation, particularly against *E. coli* biofilms. Key enzymes such as glycoside hydrolase family proteins that might have direct actions on biofilm components. Glycoside hydrolase family proteins target the polysaccharide matrix of the biofilm, hydrolyzing glycosidic bonds and leading to the breakdown of the biofilm’s structural framework.

The genomic analysis of *Escherichia* phage AUBRB02 revealed significant insights into its genetic composition and evolutionary relationships. Phylogenetic analysis (Figure 7B) placed AUBRB02 within the *Straboviridae* family, closely related to other *Pseudomonadota* phages. AUBRB02’s lack of temperate or lysogenic life cycle genes, along with the absence of virulence and antibiotic resistance genes, confirms its strictly lytic nature, making it a safe candidate for therapeutic applications.

Over all, the results of our characterization studies demonstrate that AUBRB02 is a virulent lytic phage belonging to the *Tequatrovirus* genus and clustering with *Straboviridae* phages. Whole genome sequencing confirmed the absence of virulence and antibiotic resistance genes. This phage exhibits high lytic activity against several tested *E. coli* clinical strains, with a short latent period, large burst size, and high stability across a wide range of thermal and pH conditions. To our knowledge, this is the first instance of an *E. coli* phage being isolated and characterized in Lebanon. AUBRB02 shows great promise as a candidate for phage therapy, particularly in cases where antibiotics are ineffective. Future research should validate the functions of its hypothetical proteins and evaluate AUBRB02’s efficacy in vivo to develop effective phage therapy protocols against antibiotic-resistant infections.

## MATERIALS AND METHODS

### Bacterial Strains and Growth Conditions

Bacterial strains used in this study are listed in **Table 1**. *E. coli* strains were stored at −20°C in 60% glycerol and cultivated at 37°C, with shaking at 160 rpm, in liquid Luria–Bertani broth (LB) or plated overnight at 37°C on solid LB medium with 1.5% agar. Information on antibiotic resistance of these 18 clinical strains were obtained using Broth Microdilution and Disk Diffusion Figure 2.

**Table 1:** Bacterial strains used in our study.

| Bacterial strain | Source and other characteristics |
| --- | --- |
| ***E. coli* Laboratory Strains** |  |
| ATCC® 25922™ | Serotype 06, Biotype 01 |
| ***E. coli* Clinical Strains** |  |
| 1146 | Urine |
| 1150 | Wound |
| 1164 | Urine |
| 1166 | Urine |
| 242 | Urine |
| 448 | Urine |
| 458 | Urine |
| 167 | DTA |
| 168 | Sputum |
| 169 | Sputum |
| 173 | Urine |
| 174 | Perianal |
| 177 | Perianal |
| 178 | Perianal |
| 179 | Perianal |
| 180 | Perianal |
| 182 | Perianal |
| 183 | Perianal |

### Sewage sample collection

Sewage samples were collected in a 50 ml flacon tube from Ramlet El-Bayda untreated sewage source in Beirut. Samples were placed at room temperature overnight to sediment all the debris. Samples were then centrifuged at 5000 rpm for 15 min, and the obtained supernatant was filtered using a 0.22 μm pore-size disposable syringe filter, to eliminate the remaining residues.

### Bacteriophage isolation

*E. coli* ATCC 25922 was steaked overnight at 37°C on LB agar. Sewage samples were then subjected to a round of enrichment using the previously streaked *E. coli* ATCC 25922. The obtained enriched samples were further supplemented with 10 mM CaCl_2_ to facilitate bacteriophages growth, before incubating them overnight at 37°C with shaking [32]. Enriched samples were then centrifuged at 5000 rpm for 15 min, and the obtained supernatant was filtered using a 0.22 μm pore-size disposable syringe filter, to eliminate the remaining bacterial residues. The obtained phage lysate was subjected to DLA to examine the presence of phages against *E. coli* ATCC 25922 [33]. The obtained phages appearing in the form of plaques were picked using a sterile loop and stored at 4°C in SM buffer for further investigation.

### Bacteriophage purification and enrichment

A plaque was gently picked using a disposable loop and placed in 10 ml LB broth. 0.5 McFarland *E. coli* ATCC 25922 was added into the LB broth. The enriched phage was left to incubate overnight at 37°C with shaking, then centrifuged at 5000 rpm for 15 min and filtered using a 0.22 μm pore-size disposable syringe filter. A hundred-fold serial dilution of the phage lysate was prepared and plated using the DLA technique. The following procedure was repeated for 3 times or until observing a unique morphology throughout my plate [34]. The number of plaques obtained within the most countable plate were counted to determine the Plaque Forming Unit (PFU)/ml of each phage. The obtained PFU/ml was further used to determine the Multiplicity of Infection (MOI) for each phage.

### Host range assay

Spot assay technique was conducted to determine the phage host range. Briefly ten-fold serially diluted phages were introduced in the form of spots (10 µL each) on top of the LB agar harvesting the bacterial lawn we aim to inspect. The obtained plates were incubated overnight at 37°C to detect any lysis appearance.

### One-step growth curve

The one-step growth curve was generated using [35] previous protocol with minor modifications.

*E. coli* ATCC 25922 was cultured in 990 µL LB broth to reach a 0.5 McFarland. Bacterial cells were then mixed with 0.1 µL of phages lysate with MOI 0.01. The mixture was left to incubate at room temperature for 5 min before centrifuging it at 14,000 rpm for 1 min. After centrifugation, the supernatant with free phage was removed and the pellet was suspended in 1 mL LB broth. The resuspended pellet was then incubated at room temperature, and the phage titer was then determined at an interval of 5 min for 1 hour using the DLA method.

### Adsorption rate assay

To determine the time required for AUBRB 02 to attach to its host *E. coli* ATCC *25922*, an adsorption assay was performed according to the protocol mentioned [36] with minor modifications. Briefly, 1 ml of phage lysate at MOI 0.01 was mixed with 9 ml 0.5 McFarland *E. coli* ATCC 25922. Samples were taken at different time intervals such as 0, 5, 10, 15, 20 and 25 min, and centrifuged at 12,000× *g* for 3 min. The supernatants were used for DLA assays to determine the titers of non-adsorbed phages. DLA assay was conducted in triplicates to determine AUBRB 02 PFU/ml after an overnight incubation at 37°C.

### Bacteriolytic activity assay

The lytic activity was determined using the following protocol [37] against *E. coli* ATCC 25922. *E. coli* ATCC 25922 with 0.5 McFarland was mixed in a 96 well plate with each phage separately at four different MOI’s (10, 1, 0.1, and 0.01) creating a total volume of 170 µL per each well. We have run AUBRB 02-bacteria in four replicates. In addition, we ran both a negative control having LB broth only, and a positive control harboring both the bacterial suspension at 0.5 McFarland and LB broth. The absorbance at OD 600 nm of each well was recorded over 13 hours of incubation at 37°C while shaking.

### Phage pH and thermal stability

AUBRB 02 stability was tested against a wide range of pH’s (2-13). Briefly, phage lysates were suspended in adjusted LB broth pH, to yield a pH range of 2-13, then incubated at 37°C for 2 hours. Similarly, DLA assay was performed in triplicates to determine AUBRB 02 PFU/ml after an overnight incubation at 37°C.

For thermal stability, phage lysates with pH 7 were incubated at 4, 22, 28, 37, 50, 60, 70 and 80 for 2 hours in a dry block thermostat. DLA assay was conducted in triplicates to determine AUBRB 02 PFU/ml after an overnight incubation at 37°C.

### Biofilm elimination assay

The following experiment was executed in a 96-well plate against *E. coli* ATCC 25922 biofilms. 100-fold dilution of the 0.5 McFarland *E. coli* ATCC 25922 was performed. Out of the 100-fold diluted 0.5 McFarland *E. coli* ATCC 25922, 150 µL was transferred into all wells within the 96 well plate except in the negative control well which will contain only LB broth. Within this assay we examined the effect of AUBRB 02, and Sal INF 122131 M phage (against *Salmonella infantis*), against *E. coli* ATCC 25922 biofilms. The plate carrying the bacterial suspension was incubated overnight at 28°C without shaking. Planktonic cells were removed, and the plate was washed with 1 x PBS before adding the treatments. The plate was then incubated with treatment at 28°C for 5 hours and was again washed with 1 x PBS and stained with crystal violet dye. The plate was washed again, and 95 % Ethanol was then added to detach the cells. The plate optical density of the plate content was examined at 595 nm.

### DNA Extraction

Phage DNA was extracted using the Phenol Chloroform Isoamyl alcohol protocol (Center for Phage Technology, 2018) with brief changes. Phage lysate was treated with 20 mg/ml DNase I and 10 mg/ml RNase A for 1.5 hours at 37°C. DNase I and RNase A were then deactivated with 0.5 M EDTA at 75°C for 15 min. Then 20 mg/ml Proteinase K was used at 56°C to digest the protein capsid followed by 10 % SDS. Phenol: Chloroform: Isoamyl alcohol 25:24:1 was added and centrifuged for 10 min at 10,000 xg. The supernatant was treated with Chloroform: Isoamyl alcohol 24:1 and centrifuged for 10 min at 10,000 xg. Supernatant was removed and 1/10 volume of 3 M NaOAc (pH 7.5), and 2.5 volumes of ice-cold ethanol were added. Followed by centrifuge and pellet washes using 70 % Ethanol. DNA concentration was detected using a Nanodrop.

### Phage Genome Sequencing and Bioinformatics Analysis

Genomic DNA was extracted from high-titer phage lysates. Phage DNA libraries were prepared using the Nextera kit (Illumina) and sequenced with 2 × 250 bp reads on a MiSeq system (Illumina) at the DNA Sequencing Facility of the American University of Beirut. Genome assembly and annotation of the *Escherichia coli* phage AUBRB02 was performed in a Linux environment using command-line tools. Low-quality reads were filtered, and adapters were trimmed using Trimmomatic (v0.4) [38]. The filtered reads were assembled using SPAdes (v3.15.5) [39], resulting in phage genomes resolved as single contigs with an average depth exceeding 300×. Coding sequences and tRNA genes were predicted and annotated using Prokka. Circular visualizations of the phage genomes were generated using Proksee (formerly CGView Server), a web-based tool for creating high-quality circular genome maps [40]. Further genomic analysis of phage AUBRB02 was conducted using Phage scope [41], a bioinformatics tool designed to predict phage lifecycle characteristics. This tool assesses the presence of genes associated with lysogeny, virulence, and antibiotic resistance. Phage taxonomy was determined using BLASTn against the NCBI database, and phylogenomic distances were scored using VIRIDIC (Virus Intergenomic Distance Calculator), with phage genomes showing ≥95% identity and coverage considered the same species [42]. A phylogenetic tree based on genome sequence similarities was constructed using VIPtree (v4.0) [43]. VIPtree employs the Genome-BLAST Distance Phylogeny (GBDP) method to calculate intergenomic distances. The GBDP method uses BLAST to compare genomes, calculating distances based on the number and length of matching fragments. Specifically, VIPtree uses BLASTn for nucleotide-level alignments. The distance matrix generated by GBDP was used to construct a phylogenetic tree with the FastME algorithm, which builds trees based on balanced minimum evolution criteria, ensuring accurate and robust phylogenetic relationships. A comparative map of AUBRB02 and the closely related phage was also created using VIPtree.

### Bacterial Pangenome Analysis

Bacterial genomes analyzed in this study were obtained from pathogenic bacteria identified and stored in the biorepository of the bacteriology lab at the AUB Medical Center. A total of 18 genomes were included in the pangenome analysis, conducted using Roary (v3.11.2) [44]. A pangenome phylogenetic tree was constructed based on the core genome alignment produced by Roary, which includes genes present in all analyzed strains. The resulting phylogenetic tree was visualized using iTOL (v6.0) and annotated with relevant metadata, including strain names and antibiotic resistance profiles [45].

### Statistical Analysis

Analysis and figure generation were conducted using Python. The one-way Analysis of Variance test was employed to determine if there were statistically significant differences in biofilm biomass between the phage-treated and control groups. A p-value of less than 0.05 was considered statistically significant.

### Raw Data Availability

The raw data for the *Escherichia coli* phage AUBRB02 have been deposited in publicly accessible databases. The genomic sequence is available in the European Nucleotide Archive (ENA) under the accession number ERZ22147097. The associated biosample information can be found with the accession number SAMEA114786372. Additionally, the project has been registered under the accession number PRJEB70649.

## ACKNOWLEDGMENT

T.A.A, S.E.H, A.A.F., S.K., G.M., and E.S contributed to the write-up of this review. All authors have read and agreed to the published version of the manuscript.

This research received no external funding.

All authors have read and agreed to the published version of the manuscript.

